# Screening of a chemogenetic library unveils dual-stage antimalarials with low cytotoxicity

**DOI:** 10.1101/2025.10.26.684713

**Authors:** Laura Torres, Matthew E. Fishbaugher, Melanie Lam, Alexander T. Chao, Calla Martyn, Luying Pei, Debapriya Sengupta, Maxime Dauphinais, Jonathan E. Gable, Srinivasa P. S. Rao, Steffen Renner, W. Armand Guiguemde, Sebastian A. Mikolajczak, Anke Harupa, Gabriel Mitchell

## Abstract

The global effort to eradicate malaria requires the development of drugs with novel mechanisms of action and differentiated pharmacological profiles. Both liver and blood phases of *Plasmodium* infection are essential for disease progression and transmission, making them primary targets for therapeutic intervention. In this study, we established a dual-luciferase reporter assay to simultaneously assess *Plasmodium berghei* liver-stage development and host cell viability. Screening a library of bioactive small molecules with annotated human targets revealed several inhibitors with potent liver-stage activity and minimal cytotoxicity. Further profiling against *P. falciparum* asexual blood stages identified compounds with dual-stage activity, underscoring their potential utility for development as either prophylactic or therapeutic agents. Knowledge of their human targets will facilitate investigation into host- or parasite-directed antimalarial mechanisms, thereby accelerating both biological insight and rational drug development. Our findings constitute a valuable resource for research on malaria biology and the discovery of next-generation antimalarials.

## INTRODUCTION

Malaria remains a global health challenge, affecting hundreds of millions annually and causing life-threatening complications, particularly in young children, pregnant women, travelers, and individuals with HIV (1). Most malaria cases are attributed to infection with *Plasmodium falciparum*, predominantly found in sub-Saharan Africa (1). However, *Plasmodium vivax* also accounts for a significant portion of malaria cases, exhibits a more widespread distribution, and causes clinical relapses (1, 2). *Plasmodium* parasites have repeatedly demonstrated their adaptability to environmental challenges and, despite the deployment of resources for antimalarial discovery, novel therapeutics are still required to face the emergence of drug resistance and support the global effort to eradicate malaria (3).

The complex lifecycle of *Plasmodium* provides several opportunities for therapeutic intervention (3, 4). Transmission begins when *Plasmodium* sporozoites are injected into humans by *Anopheles* mosquitoes. Shortly after the bite, sporozoites infiltrate the liver, where they invade hepatocytes and establish a parasitophorous vacuole, a specialized compartment that supports intracellular development by leveraging both host and parasite biological processes (5). Within this vacuole, parasites differentiate and replicate extensively via schizogony (5). In addition to forming replicative liver-stage schizonts, certain species such as *P. vivax* can produce hypnozoites. Hypnozoites are quiescent forms that persist for weeks to months before activating, leading to clinical relapses and continued transmission (2). Upon completion of liver-stage development, thousands of merozoites are released into the bloodstream, where they invade red blood cells and initiate the asexual replication cycle responsible for the symptomatic phase of malaria. A subset of these asexual blood-stage parasites differentiates into gametocytes, which can be taken up by mosquitoes during a blood meal and complete sexual reproduction in the mosquito midgut (3, 6).

Although the liver phase is clinically silent, it is a critical bottleneck in the parasite lifecycle and an attractive target for chemoprevention. Liver-stage inhibitors can block the establishment of blood-stage infection, prevent symptom onset, and reduce transmission. Moreover, because parasite burden is low at this stage, liver-targeting compounds may carry a reduced risk of resistance development. Dual-stage inhibitors, which are active against both liver and asexual blood stages, offer additional flexibility, as they may be developed either as preventive agents or as treatments, depending on clinical needs and resistance management strategies (3, 4).

Phenotypic screening of compound libraries against *Plasmodium* parasites has been instrumental in antimalarial drug discovery (4). However, the largest and most comprehensive screens have focused on asexual blood stages (7–9), potentially biasing hit identification toward compounds that target blood stage-specific pathways (10–12). These screens typically involve testing large numbers of structurally diverse compounds, with only a minor fraction selected for subsequent medicinal chemistry optimization to improve potency, selectivity, and pharmacological properties, ultimately yielding molecules with robust in vivo efficacy (4, 13). Elucidating the mechanism of action for hits from phenotypic screens can be challenging, especially for compounds with a high barrier to resistance (14). However, once validated, biological targets can guide more focused drug discovery, a process that can be further accelerated through computational and structure-based approaches (4, 13).

In this study, we developed a dual-luciferase assay to rapidly assess both cytotoxicity and inhibitory activity of compounds against the liver stages of *P. berghei*, a model organism that causes malaria in rodents and readily infects human hepatoma cells (5, 15). Using this assay, we screened a library of bioactive compounds, including molecules with known mechanisms of action in human cells, and identified several hits with activity against *P. berghei* liver stages and minimal cytotoxicity. Some hits also inhibited *P. falciparum* asexual blood stages, suggesting either direct antiparasitic activity or the targeting of host pathways common to both hepatocytes and red blood cells. Other hits were only active against *P. berghei* liver stages, potentially acting on liver stage-specific host or parasite pathways. Overall, this study provides detailed information on novel compounds with antimalarial activity, including chemical structures and known human targets. These compounds represent promising starting points for the development of next-generation antimalarials.

## MATERIALS AND METHODS

### Cell culture

The human hepatoma Huh7 cell line (Japanese Collection of Research Bioresources (JCRB) Cell Bank) was grown in Dulbecco’s Modified Eagle’s Medium (DMEM) with high glucose (Gibco) supplemented with 10% heat-inactivated fetal bovine serum (FBS, Corning), 100 U/mL penicillin-streptomycin (Gibco) and 1% GlutaMAX^TM^ (Gibco) (hereafter refer to as Huh7 medium) at 37 °C and 5% CO_2_.

### Generation of renilla luciferase-expressing Huh7 cells

Stable renilla luciferase-expressing Huh7 cells (Huh7_Rluc_) were generated with lentivirus from BPS Bioscience (79565-G). Briefly, cells were transduced with a multiplicity of infection (MOI) of 2 in Huh7 medium containing 8 µg/mL of polybrene (AmericanBio) followed by centrifugation at 800 × g for 1 h at 37 °C, with low acceleration and break. One day post-transduction, lentiviruses were removed, and fresh cell culture media was added to cells. Transduced cells were then selected with 250 µg/mL of geneticin (G418) for 5 weeks.

### *P. berghei* liver-stage infection assay

One day prior to infection, WT Huh7 or Huh7_Rluc_ cells were seeded at a density of 8,000 cells per well in 384-well white solid bottom plate (Greiner, 781080). *Anopheles stephensi* mosquitoes infected with luciferase-expressing *P. berghei* ANKA strain GFP-Luc_ama1-eef1a_ (*Pb*_Fluc_) (16) were produced by the SporoCore (University of Georgia, UGA) and sporozoites were isolated by microdissection of salivary glands, as previously described with modifications (17, 18). Unless otherwise specified, cells were infected with 2,000 sporozoites per well in Roswell Park Memorial Institute 1640 (RPMI, Gibco) medium containing 20% FBS (Corning). Infection was facilitated by centrifugation at 330 × *g* for 3 min at room temperature (RT), using low acceleration and braking settings. Cells were incubated with sporozoites for 2 h at 37 °C and 5% CO. The culture medium was then replaced with Huh7 medium, and compounds were added prior to further incubation for 48 hours under the same conditions.

### Dual-luciferase reporter assay

The Dual-Glo® luciferase assay (Promega) was performed as specified by the provider with modifications. Briefly, media from cell cultures was replaced with 20 µL of Dual-Glo^®^ luciferase reagent per well. Plates were incubated at RT for 10 min in the dark and shaken prior to measuring firefly luciferase (Fluc) luminescence signal. The Stop & Glo^®^ buffer was then mixed 1:1 with DMEM without phenol red (Gibco) and 40 µL was added per well. Plates were then incubated at RT in the dark for 20 min and shaken prior to measuring the renilla luciferase (Rluc) luminescence signal. Luminescence signals were read using CLARIOstar^®^ (BMG Labtech) or PHERAstar^®^ FSX (BMG Labtech) plate readers.

### Rluc assay

The Renilla-Glo^®^ luciferase assay system (Promega) was used as specified by the provider with modifications. Briefly, Huh7_Rluc_ cells were seeded at different concentrations in 384-well white solid bottom plate (Greiner) and incubated 1 day at 37°C and 5% CO_2_. The cell culture media was then removed and replaced with 10 µL of Dulbecco’s phosphate-buffer saline (D-PBS, Gibco) and 10 µL of Renilla-Glo^®^ per well. Plates were shaken, incubated at RT in the dark for 10 minutes, and the luminescence signal was read using a CLARIOstar^®^ plate reader.

### ATP cell viability assay

Promega’s CellTiter-Glo^®^ 2.0 assay kit was used as specified by the provider with modifications. Briefly, 8,000 Huh7_Rluc_ cells per well were seeded in 384-well white solid bottom plate (Greiner) and incubated 1 day at 37°C and 5% CO_2_. Cells were then treated with compounds for two days, the cell culture media was replaced with 20 µL of CellTiter-Glo^®^ reagent per well, plates were shaken and incubated at RT in the dark for 10 min, and the luminescence signal was read using a CLARIOstar^®^ plate reader.

### Screening of compounds against *P. berghei* using the dual-luciferase reporter assay

At two hours post-sporozoite infection, cell culture media was replaced, and samples were treated with a library of 6044 compounds at a final concentration of 10 μM using the Echo 650 Acoustic Liquid Handler (Beckman). This library comprises bioactive compounds with known human gene targets, including some compounds from the previously published Novartis Moa Box collection (19). Some but not all the information linked to this library is publicly accessible. Two days post-treatment, the dual-luciferase reporter assay was performed and luminescence signals were measured. Robust Z’ (RZ’) were calculated for each plates using luminescence signal values from DMSO-treated infected wells and DMSO-treated uninfected (Fluc) or empty (Rluc) wells. The data from plates with one or more RZ’ below 0.2 were excluded. Percent activities were determined by normalizing the values of each well to DMSO-treated infected wells (0% inhibition) and DMSO-treated uninfected or empty wells (100% inhibition for Fluc and Rluc, respectively) and averaged for two to four independent experiments. For hit determination, the median values of DMSO-treated infected wells minus 4× (Fluc, threshold = -84.24%) and 3× (Rluc, threshold = -41.64%) the robust standard deviations were applied as thresholds. Out of the 198 hits, 197 were further characterized - one compound was no longer available in the Novartis internal archive.

### *P. falciparum* asexual blood-stage assay

*P. falciparum* 3D7 (BEI Resources, MRA-102) parasite cultures were grown with complete medium (RPMI 1640 medium (10.4 g/L) with 0.5% AlbuMAX II, 200 μM hypoxanthine, 50 mg/L gentamicin sulfate, 35 mM HEPES, 2.0 g/L sodium bicarbonate, and 11 mM glucose) and human erythrocytes at 37°C and 5% CO_2_, as previously described (14). The antimalarial activity of compounds was measured according to a modified SYBR Green cell proliferation assay, as previously described (14).

### Structural comparison of human ATR and *Plasmodium* proteins

A structural search was conducted to identify proteins similar to human ATR (UniProt ID Q13535; PDB ID 5YZ0) among all *P. falciparum* proteins using Foldseek (20) incorporated into UniProt (search parameters = AlphaFold/UniProt50v4 + PDB100 20240101, mode: 3Di/AA, taxonomic filter: *Plasmodium falciparum*, iterative search off). The top *P. falciparum* hit, an AlphaFold model for a phosphatidylinositol 4-kinase (PI(4)K, UniProt ID Q8I406), showed structural alignment between residues 1280-1556 and human ATR residues 2320-2632. The structural similarity between *Plasmodium* PI(4)K_1280-1556_ and human ATR_2320-2632_ was confirmed using PyMOL (RMSD, root mean square deviation: 1.832 Å; The PyMOL Molecular Graphics System, Version 3.1.3 Schrödinger, LLC).

### PI(4)K enzymatic assay

A PI(4)K enzymatic assay was performed as previously described (21). Briefly, the assay was performed for 1h at RT in a 384-well black plate using the Transcreener ADP_2_ FP detection kit (BellBrook) and a final assay volume of 10 µL containing 1nM of purified *P. vivax* PI(4)K protein, 10mM Tris, pH 7.5, 1mM DTT, 10 µM ATP, 5mM Mn^2+^, 0.2% Triton X-100 and 11 µM L-α-phosphatidylinositol (dissolved in 3% *n*-octylglucoside). The reaction was then stopped by adding 10 µL of detection mix containing 1 × stop buffer, 4 nM AMP Alexa Fluor 633 tracer, and 20 µg/mL of ADP antibody. Fluorescence polarization measurements were performed on the Infinite M1000 plate reader (Tecan) with λ_ex_= 635 nm and λ_em_= 680 nm (bandwidth, 20 nm).

### Compounds, dose response treatments and analyses

All compounds were obtained from the Novartis internal archive and resuspended in DMSO. For usual dose-response experiments, compounds from master plates, containing 7- to 10-points 3-fold or half-log serial dilution series, were transferred in assay plates using the mosquito^®^ HV liquid handler (SPT Labtech), with maximal concentrations of 10 μM. Results were normalized (100%, DMSO-treated samples) and plotted with curve fitting on either GraphPad Prism (version 9.5) or DAVID Helios (version 3.2.0.364). For hit compounds, IC_50_ and CC_50_ values underwent transformation prior to medians, selectivity indexes (SIs), and IC_50_ *Pf* BS / IC_50_ *Pb* LS ratios determination (see Supplementary data S1). Briefly, data outside the limits of experimental detection were imputed to half the next lower or higher untested concentration, based on the serial dilution technic used for each assay. Subsequently, data were subjected to log10-transformation and median values were calculated. To compute SIs and IC_50_ *Pf* BS / IC_50_ *Pb* LS ratios, the data were reverted to their original scale. For accuracy, median values were re-expressed along with their corresponding experimental detection limits. Numbers of technical replicates and independent experiments are indicated in Figure legends.

### Annotation of hit compounds

Data for hit compounds were gathered from ChEMBL v35 (downloaded on 2024-12-01) and PubChem (downloaded on 2025-01-22). Target scores were determined as previously described (22).

### Preparation of graphs and artworks

BioRender (https://biorender.com), GraphPad Prism (v9.5), ChemDraw Professional (v23.1.1.3) and Adobe Illustrator (v29.8.1) were used to create figures.

## RESULTS

### A dual-luciferase reporter assay for assessing *P. berghei* liver-stage development and host cell viability

A dual-luciferase assay was customized to simultaneously evaluate parasite growth and host cell viability during the liver phase of malaria infection (Fig. 1A). As a reporter of host cell viability, the renilla luciferase (Rluc) gene was introduced into Huh7 hepatoma cells via lentiviral transduction, yielding a luminescence signal that strongly correlated with cell number (Fig. S1A). These Rluc-expressing cells (Huh7_Rluc_) were then infected with *P. berghei* sporozoites expressing the firefly luciferase (Fluc) gene (*Pb*_Fluc_) (16) for a 2-day incubation period (Fig. 1A, step 1). Parasite and host signals were then quantified using the Dual-Glo^®^ luciferase assay, which enables sequential measurement of Fluc (Fig. 1A, step 2) and Rluc (Fig. 1A, step 3) luminescence. Both signals were elevated by over two orders of magnitude compared to uninfected or untransduced controls (Fig. 1B-C). Moreover, the Fluc signal exhibited a strong positive correlation with the number of *Pb*_Fluc_ sporozoites used for infection (Fig. S1B), demonstrating the assay’s sensitivity to parasite load.

**Figure 1.**
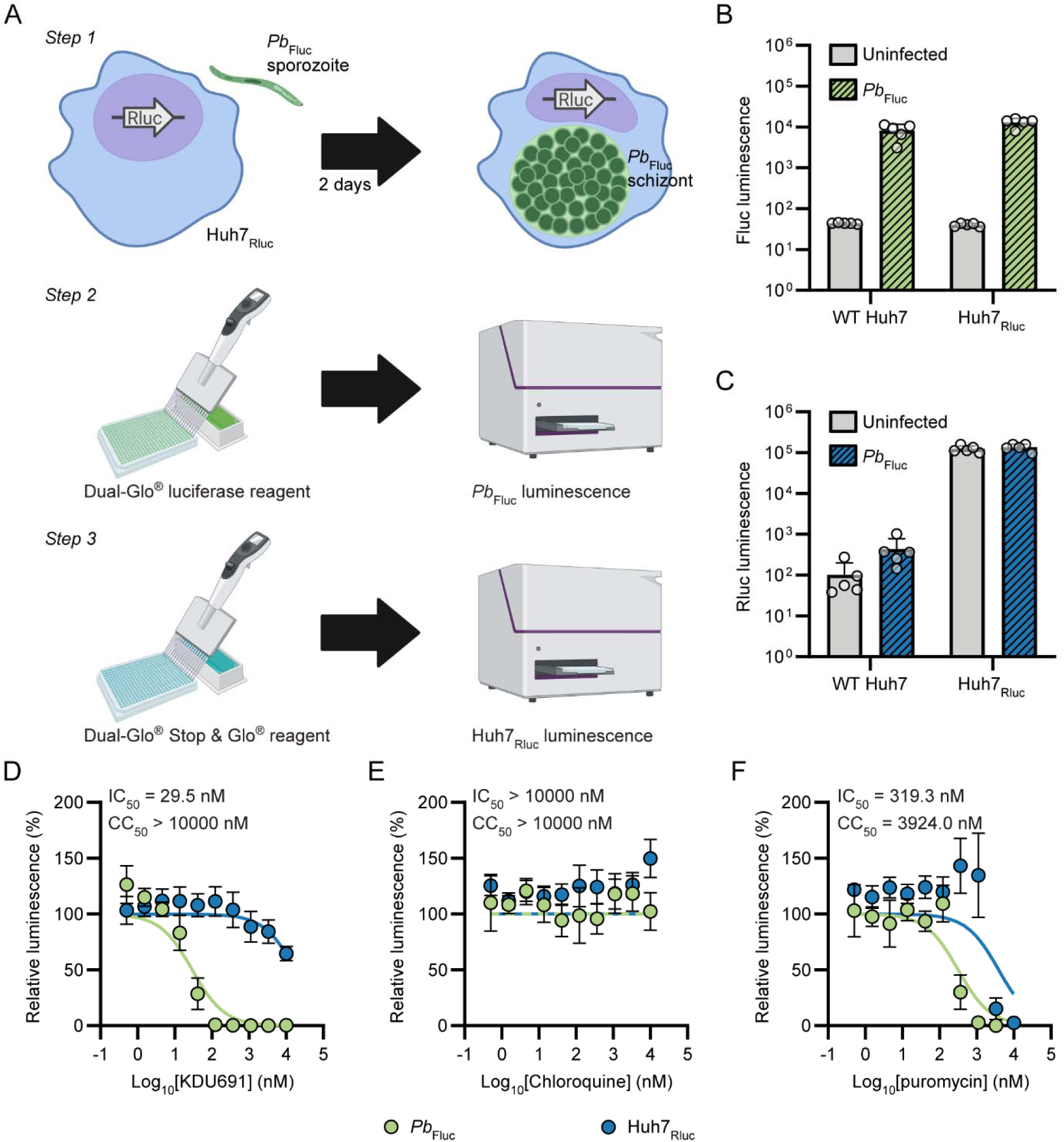
A dual-luciferase reporter assay for simultaneous assessment of *P. berghei* liver-stage development and host cells viability. (A) Schematic overview of the dual-luciferase reporter assay. (Step 1) Renilla luciferase (Rluc)-expressing Huh7 (Huh7_Rluc_) cells were infected with firefly luciferase (Fluc)-expressing *P. berghei* (*Pb*_Fluc_) and incubated for 2 days prior measuring (Step 2) *Pb*_Fluc_ and (Step 3) Huh7_Rluc_ luminescence. Fluc (B) and Rluc (C) luminescence of WT Huh7 and Huh7_Renilla_ cells infected with *Pb*_Fluc_ for 2 days. Results are unnormalized and presented as means and SDs from one experiment with 5 technical replicates (n=5). Dose-response effect of the liver-stage inhibitor KDU691 (D), the blood-stage inhibitor chloroquine (E), and the cytotoxic control puromycin (F) on Fluc and Rluc luminescence signals. Half-maximal inhibitory concentration (IC_50_) and half-maximal cytotoxic concentration (CC_50_) values are shown for each compound. Data are normalized and expressed as means and SEMs from 4 independent experiments (N=4, with technical duplicates). 100% corresponds to DMSO-treated infected samples.

To confirm the suitability of this assay for compound screening during liver-stage infection, we first tested three reference compounds: KDU691 (Fig. 1D), a known inhibitor of liver stages (21); chloroquine (Fig. 1E), which is active against asexual blood stages but not liver stages (23); and puromycin (Fig. 1F), a cytotoxic control compound. As anticipated, KDU691 reduced the *Pb*_Fluc_ signal with a half-maximal inhibitory concentration (IC_50_) of 16.0 - 29.5 nM, consistent with previously reported values (21, 24), and showed no cytotoxicity toward Huh7_Rluc_ cells, with a half-maximal cytotoxic concentration (CC_50_) greater than 10,000 nM (Fig. 1D and S2D). Also in line with expectations, chloroquine did not reduce either luminescence signal (Fig. 1E and S2F) (24), while puromycin affected both parasite and host cell signals (Fig. 1F and S2G) (see GNF-*Pf*-2016 in (25)). It is noteworthy that puromycin impacted *Pb*_Fluc_ at lower concentrations (IC_50_ = 231.0 - 319.3 nM) than Huh7_Rluc_ (CC_50_ = 2994 -3924 nM), possibly due to the faster replication rate of liver-stage parasites compared to that of host cells. Potency against *Pb*_Fluc_ was also in agreement with previously reported data for other established antimalarials, including cabamiquine (24, 26), atovaquone (24, 27, 28) and KAF179 (11) (Fig. S2). KAE609 showed only weak activity against *P. berghei* liver stages (IC_50_ = 4607 nM) (Fig. S2), consistent with earlier findings (11, 29) but contrasting with a recent report by Nguyen et al. (24). Notably, KAE609 showed minor cytotoxicity at higher concentrations, which may have contributed to the observed weak *Pb*_Fluc_ activity (Fig. S2). We further confirmed cell viability data for these antimalarials using an orthogonal assay (CellTiter-Glo®) that quantifies viable cells based on ATP levels (Fig. S2), and found that results were consistent across both assays. Overall, these results demonstrate that the dual-luciferase assay is a reliable and sensitive platform for evaluating compound activity against *P. berghei* liver stages and host cells.

### Screening of a bioactive library against *P. berghei* liver stages using the dual-luciferase reporter assay

The dual-luciferase reporter assay was employed to screen a library of 6044 compounds, partially overlapping with the previously published Novartis Moa Box collection (19). This library consists of bioactive compounds with annotated human targets, sourced from both public and internal datasets. Compounds were screened at a final concentration of 10 μM across 2-4 independent experiments (Fig. 2A). As a quality control measure, robust Z’ (RZ’) values were calculated for each plate and luminescence signal, and plates with RZ’ values below 0.2 were excluded from further analysis (Fig. 2B). The means and standard deviations of RZ’ for the Fluc and Rluc signals were 0.61 ± 0.21 and 0.76 ± 0.16, respectively, confirming the robustness and reliability of the assay. As expected from a bioactive collection, several compounds impacted liver-stage development and host cell viability (Fig. 2C). It is worth noting that some compounds induced an increase in the Rluc signal relative to DMSO-treated controls (Fig. 2C), likely reflecting a known technical artifact. Indeed, this effect is consistent with a previous report showing that certain compounds, including staurosporine, can enhance the activity of the CMV promoter (30), which drives Rluc expression in our system (Fig. S3). To select liver-stage inhibitors for follow-up studies, cutoffs were applied (see Materials and Methods), resulting in a set of 198 compounds with potent activity against *P. berghei* liver stages and minimal cytotoxicity (Fig. 2C). Together, these results demonstrate the utility of the dual-luciferase assay for compound screening. They also suggest that screening the bioactive library has led to the discovery of *Plasmodium* liver-stage inhibitors.

**Figure 2.**
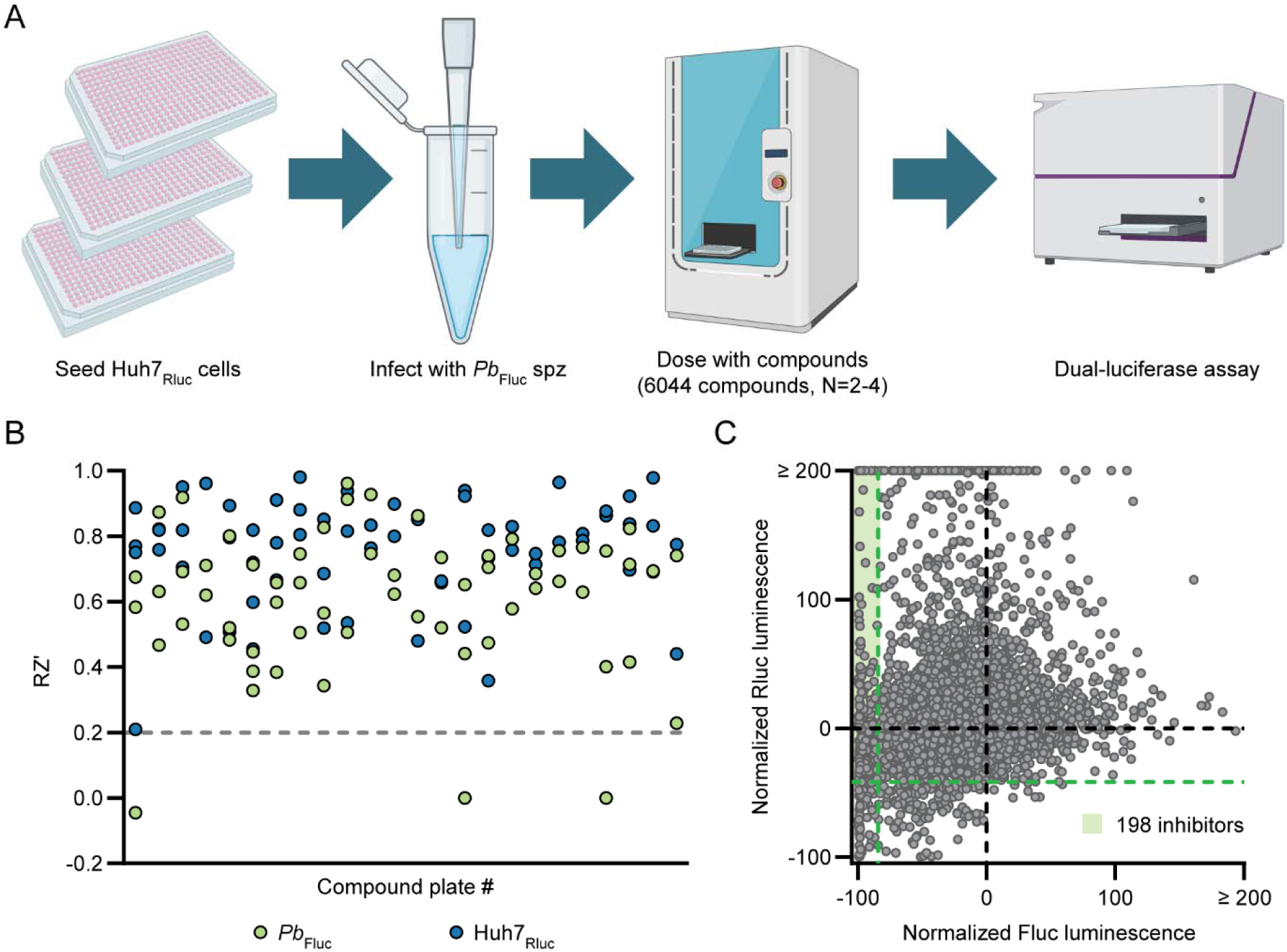
Screening of a chemogenetic library against *P. berghei* liver stages using the dual-luciferase reporter assay. (A) Diagram outlining the screening methodology. Huh7_Rluc_ cells were seeded in 384-well plates, infected with firefly luciferase-expressing *P. berghei* (*Pb*_Fluc_) sporozoites for 2h, treated with 10 μM of each compound using an acoustic liquid handler, and incubated for a total of 2 days prior to performing the dual-luciferase reporter assay. The impact of each compound was assessed across 2-4 independent experiments (N=2-4). (B) Robust Z’ (RZ’) were calculated for each plate and for both firefly luciferase (Fluc, in green) and renilla luciferase (Rluc, in blue) luminescence readouts. Plates with at least one RZ’ value below 0.2 (gray dashed line) were excluded from further analysis. (C) Fluc and Rluc luminescence values associated with each compound. Data are normalized and presented as mean values. 0% (black dashed line) and -100% correspond to no and complete inhibition, respectively. Green dashed lines indicate the Fluc and Rluc luminescence thresholds for the determination of hits, with compounds in the green shaded area selected for further characterization.

### Dose-response validation of hit compounds against *P. berghei* liver stages

Compounds identified in the single-point screen were further validated through dose-response experiments using the dual-luciferase reporter assay (Fig. 3 and Supplementary data S1). These experiments enabled the determination of *Pb*_Fluc_ IC_50_s (Fig. 3A), Huh7_Rluc_ CC_50_s (Fig. 3B) and selectivity indexes (SIs) (Fig. 3C), calculated as the ratio of CC_50_ to IC_50_, which reflects the window between cytotoxicity and antimalarial activity. As a benchmark, the cytotoxic control compound puromycin yielded an SI of 9.4, which was used as an approximate cutoff for hit prioritization. Compounds were ranked by SI and visualized in a heat map, highlighting those with potent activity against *P. berghei* liver stages (Fig. 3D). A total of 155 compounds exceeded an SI of 10 (Supplementary data S1), indicating selective antimalarial activity. For the majority of these hits, chemical structures and annotated mechanisms of action in human cells are publicly available (Supplementary data S1), which may facilitate downstream target identification (4, 7, 31, 32). Among the identified hits, several compounds have documented activity against asexual blood stages, including WR99210 (Hit # 5) (33, 34), LY-411575 (Hit # 7) (35), atovaquone (Hit # 9) (36, 37), pyrimethamine (Hit # 27) (7, 33), apicidin (Hit #32) (38), compound 3 / GNF-*Pf*-4076 (Hit #40) (25, 39), cycloguanil (Hit # 58) (7), and norketotifen (Hit # 59) (40) (Fig. 3D), suggesting that some of the newly identified compounds may possess multistage activity. Conversely, clomiphene (Hit # 47), a compound with previously reported liver stage-specific activity (32), was also among the identified hits (Fig. 3D). Overall, these results confirm that screening the bioactive library using the dual-luciferase reporter assay identified *Plasmodium* inhibitors.

**Figure 3.**
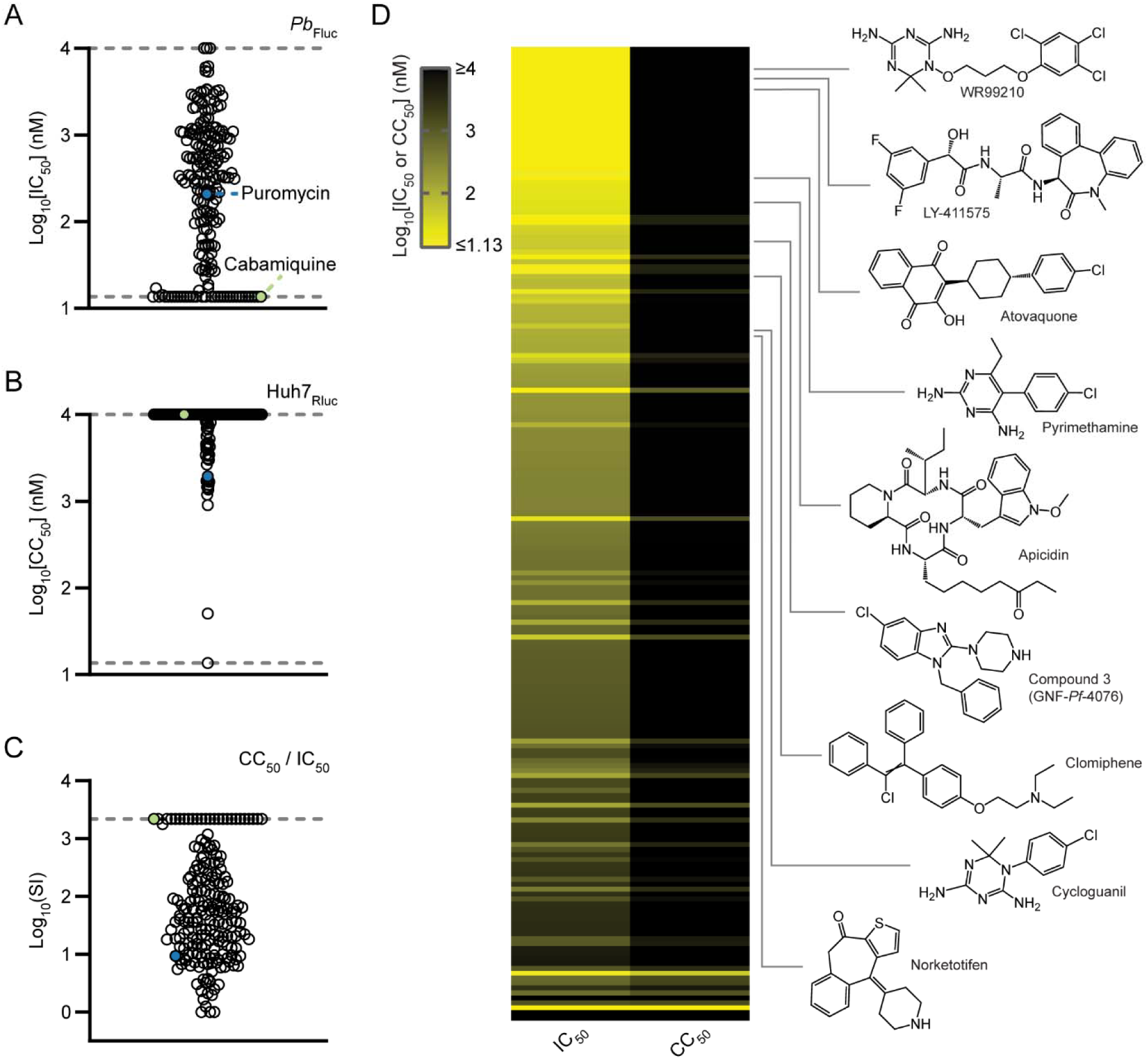
Dose-response validation of hit compounds against *P. berghei* liver stages. The dose-response activity of hit compounds was assessed against *P. berghei* liver stages at 2 days post-infection using the dual-luciferase reporter assay. Median IC_50_s (A), median CC_50_s (B) and approximative selectivity indexes (SIs) (C) were determined from 2-4 independent experiments (N=2-4). Cabamiquine (in green) and puromycin (in blue) served as antimalarial and cytotoxic controls, respectively. Limits of detection are illustrated by grey dashed lines. Values below or above the limit of detection were imputed to allow the determination of medians and SIs. (D) Heat map illustrating median IC_50_s and CC_50_s for hit compounds. Compounds are organized from top to bottom based on their SI. Molecular structures of compounds with known antimalarial activity are showcased. Some data are shared between Fig. 3-5.

### Profiling hit compounds against *P. falciparum* asexual blood stages

Given that several identified hits have known activity against *Plasmodium* asexual blood stages (Fig. 3D), an expanded set of compounds was tested to assess cross-stage efficacy (Fig. 4 and Supplementary data S1). Notably, a subset of *P. berghei* liver-stage inhibitors also demonstrated activity against *P. falciparum* blood stages (Fig. 4A), resulting in a moderate positive correlation between liver-stage and blood-stage IC_50_s (*r* = 0.5309) (Fig. 4B). Among compounds with an SI > 10, we identified 65 dual-stage (IC_50_ *Pf* BS / IC_50_ *Pb* LS < 10) and 55 potential liver stage-specific (IC_50_ *Pf* BS / IC_50_ *Pb* LS > 10) inhibitors (Fig. 4C, Supplementary data S1). These findings suggest that a substantial fraction of the inhibitors target biological pathways essential for both liver-stage and asexual blood-stage development.

**Figure 4.**
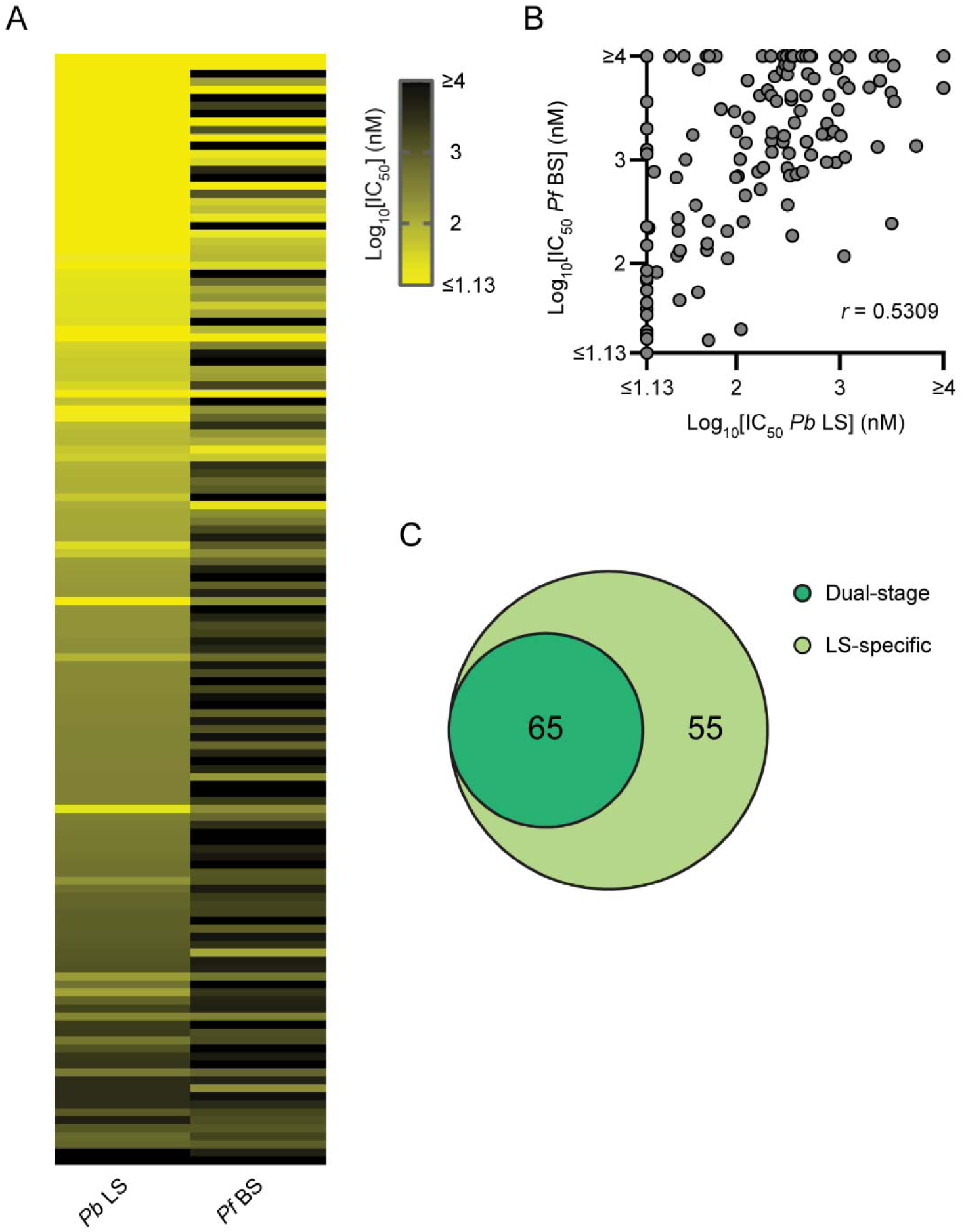
Characterization of hit compounds against *P. falciparum* asexual blood stages. (A) Heat map displaying median IC_50_s of hit compounds against *P. berghei* (*Pb*) liver stages (LS) and *P. falciparum* (*Pf*) blood stages (BS). (B) Graph showing the correlation between *P. berghei* liver-stage and *P. falciparum* blood-stage IC_50_s. Median IC_50_s against *P. falciparum* blood stages were obtained from at least one experiment with technical duplicates. The Pearson correlation coefficient (*r*) is shown. Values outside the limits of detection were imputed to calculate medians and harmonized between assays for graphical representation. (C) Venn diagram showing numbers of compounds with dual-stage (IC_50_ *Pf* BS / IC_50_ *Pb* LS < 10) and liver stage-specific (IC_50_ *Pf* BS / IC_50_ *Pb* LS > 10) antimalarial activities. Only compounds with an SI (CC_50_ Huh7 / IC_50_ *Pb* LS > 10) were considered. Some data are shared between Fig. 3-5.

Conversely, compounds that exclusively inhibit *P. berghei* liver stages may act on host or parasite pathways uniquely required during liver-stage infection. Alternatively, they may exhibit species-specific activity across different *Plasmodium* parasites or constitute technical artifacts (41). Because the liver stage-specific inhibitors identified in this study are supported solely by data from the dual-luciferase reporter assay, further validation is required to confirm their biological relevance. Nevertheless, given the complementarity between the *P. berghei* liver-stage and the *P. falciparum* blood-stage assays, this study confidently identified a collection of dual-stage inhibitors of malaria.

### Dual-stage inhibitors of *Plasmodium* parasites

The chemical structures and annotated human targets of several *Plasmodium* inhibitors highlighted in this study are publicly available (Supplementary data S1). Among the dual-stage inhibitors, notable examples include ATR inhibitors VE-822 and VE-821 (42, 43), a signal peptide peptidase-like 2A (SPPL2A) inhibitor (44), the γ-secretase and pan-Notch inhibitor Varegacestat (45), histone deacetylase (HDAC) inhibitors (Nanatinostat (46) and Hit # 31 (47)), the DNA topoisomerase I inhibitor Exatecan (48, 49), and a RabGGTase inhibitor (50, 51) (Fig. 5). While these compounds are known to act on host targets and pathways, their antimalarial activity may arise from the inhibition of *Plasmodium* homologs, particularly for validated parasite targets such as HDACs (38, 52) and DNA topoisomerases (53, 54). Moreover, knowledge of a compound’s human target can facilitate the identification of structurally related parasite proteins that may also be susceptible to inhibition. For example, we found that *P. falciparum* phosphatidylinositol 4-kinase (PI(4)K) (PF3D7_0509800) shares structural features with human ATR (Fig. S4). Consistent with this, ATR inhibitors VE-822 and VE-821 (Fig. 5) exhibit structural similarity to the known *Plasmodium* PI(4)K inhibitor MMV390048 (55) and potently inhibit recombinant *P. vivax* PI(4)K *in vitro* (Fig. S4). These findings suggest that our dual-stage inhibitors may exert antimalarial effects through parasite targets. Overall, this study has identified a diverse set of dual-stage antimalarial compounds, offering valuable starting points for target identification and therapeutic development.

**Figure 5.**
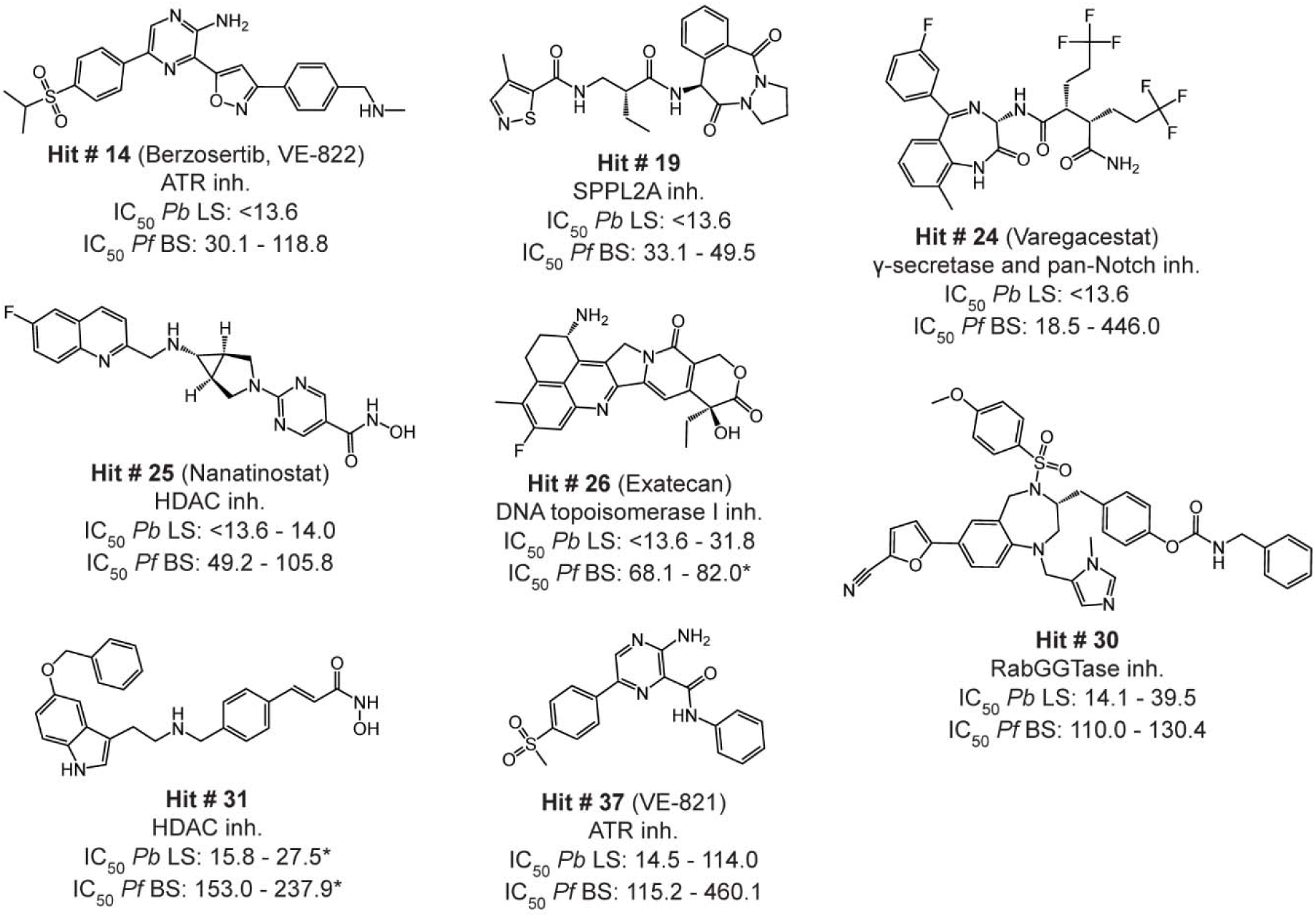
Dual-stage inhibitors of *Plasmodium* parasites. Structures of select dual-stage inhibitors are featured. Primary known mechanism of action in human and ranges of IC_50_s (i.e., min - max nM values) against *P. berghei* (*Pb*) liver stages (LS) and *P. falciparum* (*Pf*) asexual blood stages (BS) are shown. Compounds were selected based on their antimalarial activities and the absence of detectable cytotoxicity in Huh7_Rluc_ cells (CC_50_ > 10000 nM). Results are from at least two independent experiments (N=2). ATR, ataxia telangiectasia and Rad3-related; HDAC, histone deacetylase; RabGGTase, Rab geranylgeranyl-transferase; SPPL2A, signal peptide peptidase-like 2A. * indicates that one outlier value was excluded from the corresponding IC_50_ range. Some data are shared between Fig. 3-5.

## DISCUSSION

The liver phase of *Plasmodium* infection represents a critical window for therapeutic intervention against malaria, complementing approaches targeting the symptomatic blood phase of infection (3, 4). During the liver phase, parasites undergo intracellular replication within hepatocytes, exploiting both host and parasite processes (5) that may be amenable to pharmacological modulation. However, effective liver-stage therapeutics must inhibit parasite development without significant adverse effects on host cells and tissues. To expedite the discovery of non-cytotoxic antimalarials, we tailored a dual-luciferase reporter assay capable of simultaneously quantifying *Plasmodium* liver-stage development and host cell viability, enabling immediate exclusion of cytotoxic compounds (Fig. 1). Using this assay, we screened a chemogenetic library of small molecules with annotated human targets (Fig. 2 and Supplementary data S1) and identified novel inhibitors of liver-stage parasites (Fig. 3). Several of these compounds also suppressed *P. falciparum* asexual blood stages (Fig. 4 and 5), suggesting dual-stage activity.

Although chemogenetic libraries provide a powerful approach for accelerating target identification (19, 56), dissecting compound mechanisms during liver-stage infection remains challenging. Homologous proteins in host and parasite can complicate target attribution and validation, particularly when phenotypes may arise from effects on either or both. However, our compounds that inhibit both liver and blood stages are hypothesized to act on parasite targets conserved across developmental stages and species, rather than on host pathways, as red blood cells lack nuclei and most organelles and therefore offer fewer host targets (57). For instance, Hit # 30 may inhibit *Plasmodium* RabGGTase (Fig. 5), potentially disrupting Rab protein prenylation and membrane localization (58, 59), suggesting that prenyltransferases may represent a broader class of antimalarial targets (51, 60). As another example, human ATR inhibitors VE-822 and VE-821 may act on parasites by inhibiting *Plasmodium* PI(4)K (Fig. S4). These examples illustrate how annotated human targets, especially when integrated with structure-based approaches, can inform parasite target discovery and accelerate mechanistic insight.

In addition to dual-stage inhibitors, we also identified compounds with potential liver stage-specific activity. These may act on parasite pathways uniquely required for liver-stage development, on host pathways, or reflect species-specific biology, as liver-stage activity was assessed in *P. berghei* and blood-stage activity in *P. falciparum*. An example of a host-directed mechanism is the inhibition of liver stages by the cholesterol trafficking and biosynthesis inhibitor U18666A (61, 62) (Hit # 12, Supplementary data S1; (63)), consistent with the established role of host cholesterol in supporting parasite development (63, 64). Our results also suggest that liver-stage parasites may be sensitive to inhibition of host hypoxia-inducible factor (HIF)-1α (Moracin P, Hit #22; Supplementary data S1), a transcription factor mediating cellular responses to hypoxia in addition to its roles under normoxic conditions (65–67). Consistent with this finding, a previous study indicated that hypoxia promotes the development of *Plasmodium* liver stages through mechanisms partly mediated by HIF-1α (68). Interestingly, HIFs and the hepatic oxygen gradient have been implicated in the metabolic zonation of the liver (69), with the perivenous lobule zone—characterized by lower oxygen tension—being more permissive to liver-stage development (68, 70). As such, HIF-1α may contribute to the differential susceptibility of hepatocyte subpopulations to *Plasmodium* infection. Overall, these findings highlight potential host-parasite interactions that warrant further investigation.

The chemically diverse hits identified in this study broaden the antimalarial discovery space and include potent, low-toxicity inhibitors with structures and annotations that support mechanism-of-action deconvolution and medicinal chemistry optimization. Although further profiling is needed to assess their suitability, some compounds may serve as starting points for developing chemotypes with potential relevance to MMV’s long-acting chemoprotection strategy (71). In summary, this study lays the groundwork for future efforts to validate novel targets, optimize lead compounds, and inform the design of next-generation antimalarial drugs for both prevention and treatment.

## ACKNOWLEDGEMENTS

The authors thank Thomas Krucker and Thierry T. Diagana for providing their support and mentorship during the execution of this study. The authors also thank Harry Cheung, Allen Cornett, Richard T. Eastman, Erika L. Flannery, Rajiv S. Jumani, Eric J. Martin, Beat Nyfeler, Maude Patoor and Joseph M. Young for helpful discussions and technical assistance, and the SporoCore group (University of Georgia) for producing *P. berghei*-infected mosquitoes. This work was supported by a grant from the Gates Foundation (INV010720; Thierry T. Diagana). The conclusions and opinions expressed in this work are those of the authors alone and shall not be attributed to the Gates Foundation. All authors were employed by and/or shareholders of Novartis Pharma AG during this study.

## AUTHOR CONTRIBUTIONS

**Laura Torres**: Conceptualization, Data curation, Formal analysis, Investigation, Methodology, Visualization, Writing - original draft. **Matthew E. Fishbaugher**: Investigation, Methodology. **Melanie Lam**: Investigation. **Alexander T. Chao**: Investigation, Data curation, Formal analysis. **Calla Martyn**: Formal analysis, Software. **Luying Pei**: Investigation, Data curation. **Debapriya Sengupta**: Investigation, Data curation. **Maxime Dauphinais**: Formal analysis, Visualization. **Jonathan E. Gable**: Formal analysis. **Srini P.S. Rao**: Conceptualization, Supervision. **Steffen Renner**: Conceptualization, Formal analysis, Software. **W. Armand Guiguemde**: Methodology, Supervision. **Sebastian A. Mikolajczak**: Conceptualization, Supervision. **Anke Harupa**: Conceptualization, Investigation, Methodology, Supervision, Writing - original draft. **Gabriel Mitchell**: Conceptualization, Formal analysis, Investigation, Methodology, Project administration, Supervision, Visualization, Writing - original draft. **All authors**: Writing - review & editing.

## ADDITIONAL FILES

**Figure S1.**
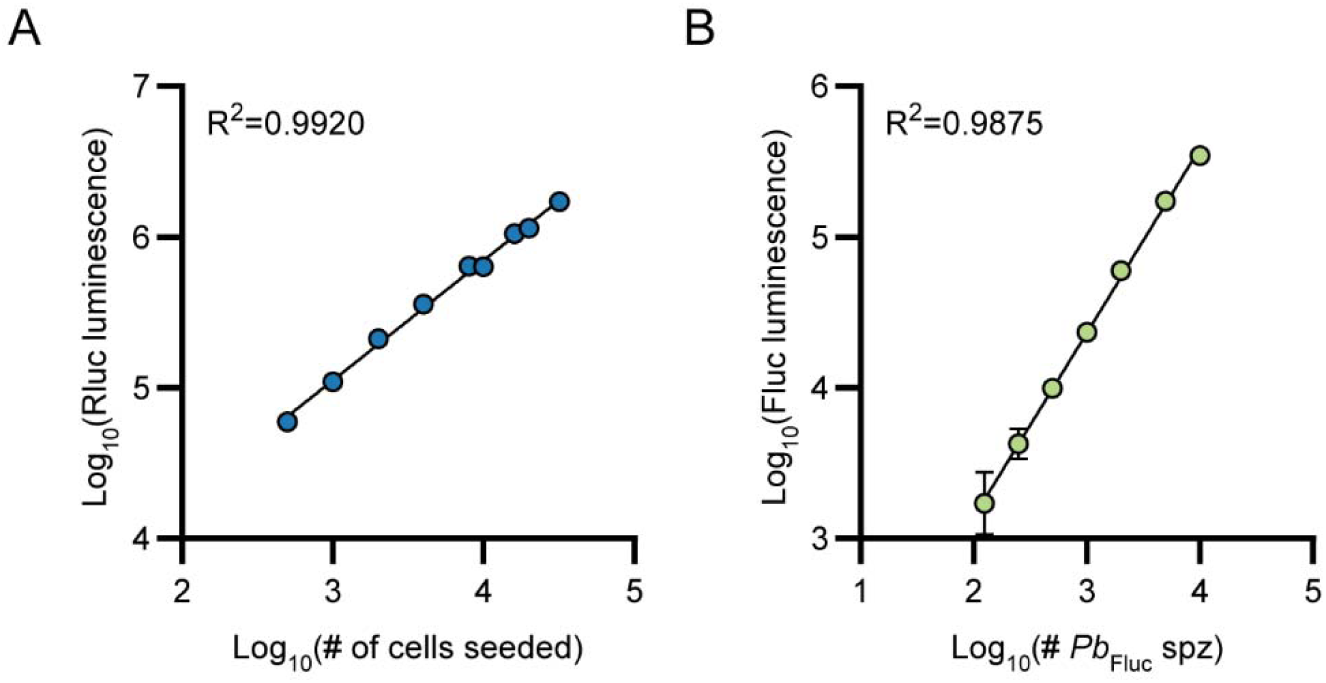
(A) Renilla luminescence (Rluc) as of function of the number of Huh7_Rluc_ cells seeded per well of a 384-well plate. Results were obtained from uninfected cells, using the Rluc assay, and are represented as means and SDs from one experiment (N=1) performed with six technical replicates. (B) Firefly luciferase (Fluc) luminescence as of function of the number of Fluc-expressing *P. berghei* (*Pb*_Fluc_) sporozoites used to infect cells for 2 days. Results were obtained using the dual-luciferase reporter assay and are represented as means and SDs from one experiment (N=1) performed with four technical replicates. Coefficients of determination (R^2^) from simple linear regressions are shown.

**Figure S2.**
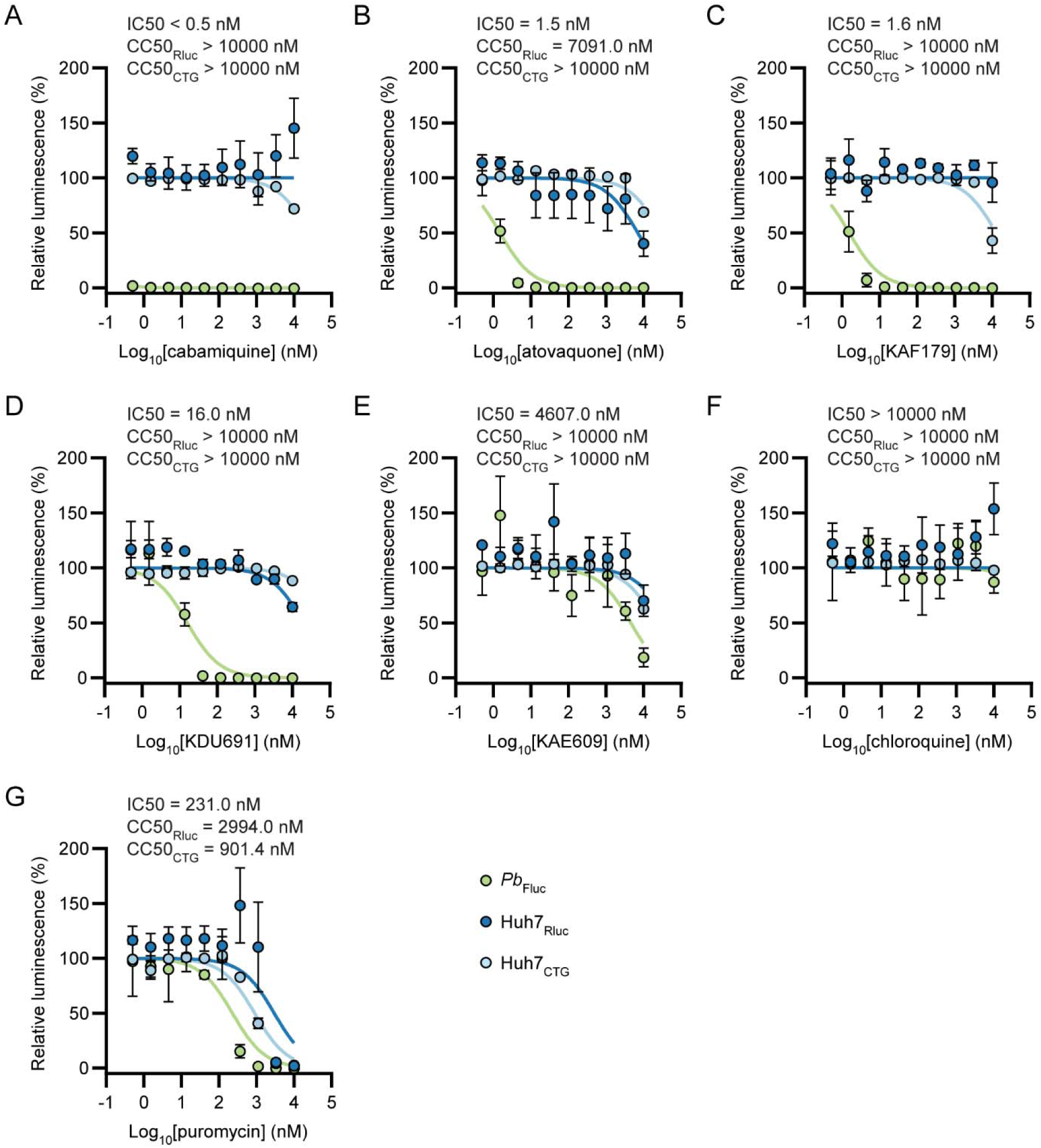
Dose-response graphs showing the impact of cabamiquine (A), atovaquone (B), KAF179 (C), KDU691 (D), KAE609 (E), chloroquine (F) and puromycin (G) on *P. berghei* (*Pb*_Fluc_) liver-stage development and Huh7 (Huh7_Rluc_) viability as determined using the dual-luciferase reporter assay 2 days post-infection. The toxicity of compounds was also assessed in uninfected cells using the CellTiter-Glo^®^ (CTG) assay. Results are presented as means and SEMs from 3-4 independent experiments (N =3-4) performed with 2-4 technical replicates. IC_50_s (*Pb*_Fluc_) and CC_50_s (Huh7_Rluc_ and Huh7_CTG_) are shown above each graph. 100% corresponds to DMSO-treated samples. Some data are shared between Fig. 1 and Fig. S2.

**Figure S3.**
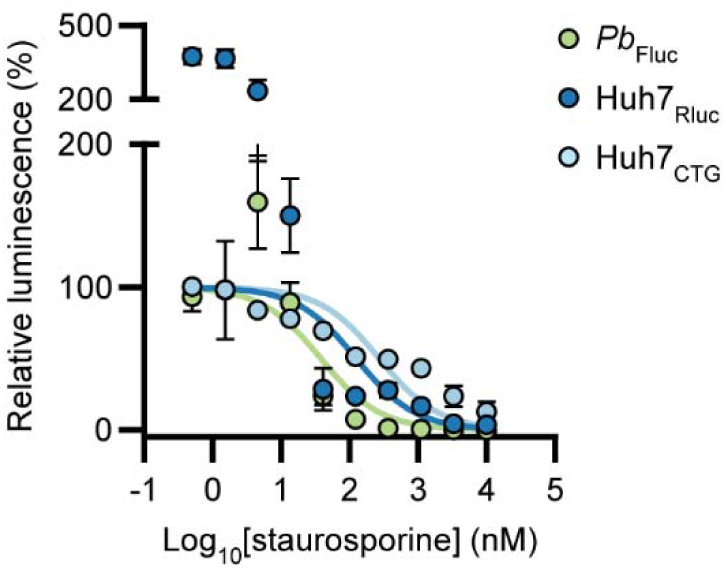
Dose-response graph showing the effect of staurosporine on *P. berghei* (*Pb*_Fluc_) liver-stage development and Huh7 (Huh7_Rluc_) viability as determined using the dual-luciferase reporter assay 2 days post-infection. The toxicity of compounds was also determined in uninfected cells using the CellTiter-Glo^®^ (CTG) assay. Results are presented as means and SEMs from 3-4 independent experiments (N=3-4) performed with technical duplicates. 100% corresponds to DMSO-treated samples.

**Figure S4.**
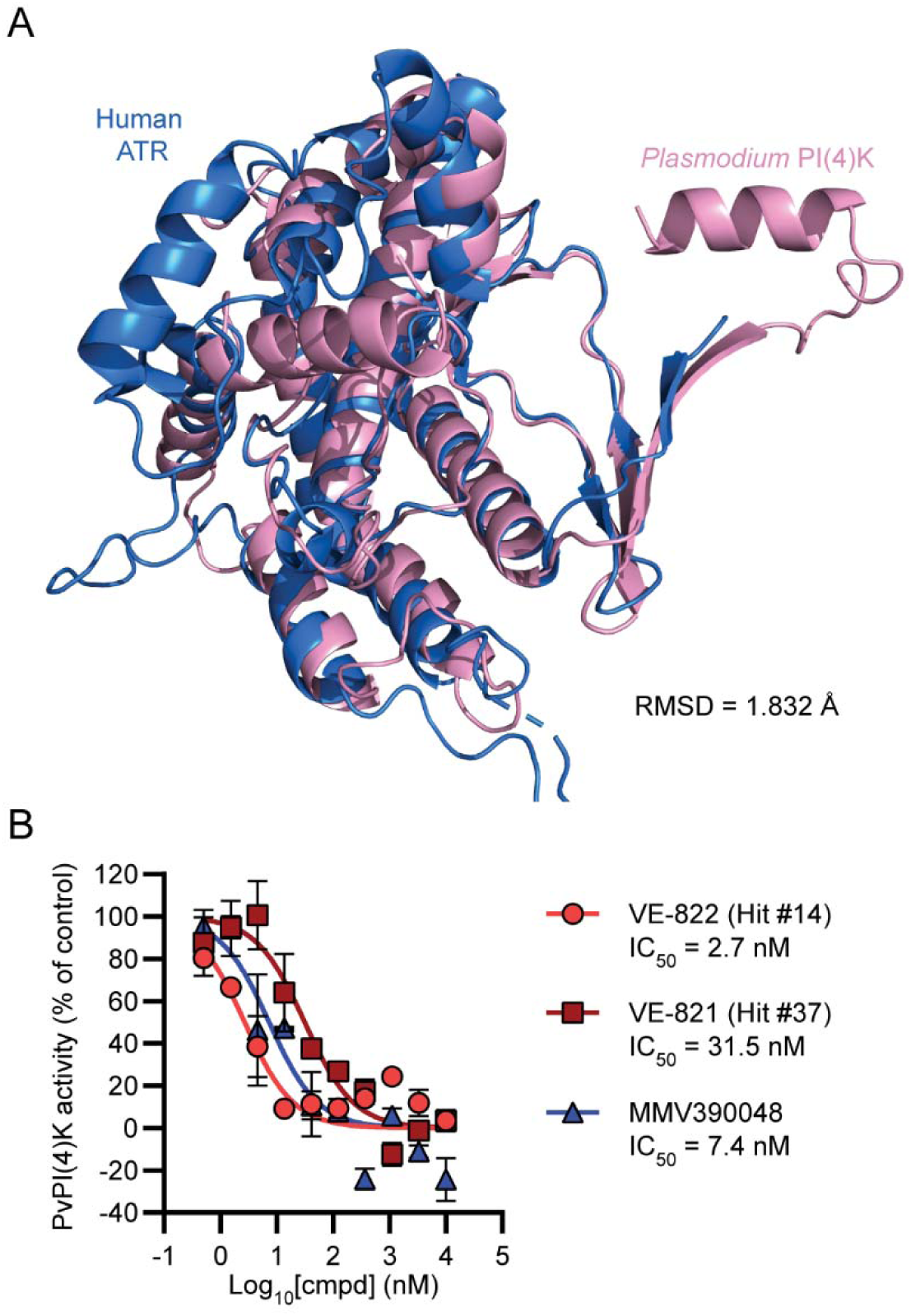
(A) Structural alignment of *P. falciparum* PI(4)K (amino acid 1280-1556, in pink) and human ATR (amino acid 2320-2632, in blue). The alignment was performed using PyMol as described in the Materials and Methods. RMSD, root mean square deviation. (B) Effect of ATR inhibitors VE-822 (Hit #14) and VE-821 (Hit #37) and a control *Plasmodium* PI(4)K inhibitor (MMV390048) on the activity of recombinant *P. vivax* (Pv) PI(4)K. Results are presented as means and SDs from one experiment (N=1) performed with technical duplicates. IC_50_s are shown. 100% corresponds to DMSO-treated samples.

**Supplementary data S1.** (Summary table sheet) InChIKey, preferred name(s), top human target, mean single-point activity (%) against *P. berghei* (*Pb*_Fluc_) liver stages (LS) and Huh7_Rluc_, median *Pb*_Fluc_ LS IC_50_s and Huh7_Rluc_ CC_50_s, SIs, and median *P. falciparum* asexual blood-stage (BS) IC_50_s for hit compounds. Only publicly available information is listed, except for the profiling of antimalarial activities. (Cmpd characteristics sheet) InChIKey, ChEMBL IDs, compound ID (CID) number and PubChem names for hit compounds. Only publicly available information is listed. (Human cmpd target sheet) Human targets, including the target with best score, for hit compounds. Only publicly available information is listed. (*Pb* LS dual luciferase) Raw and transformed data for the dose-response effect of hit compounds on *Pb*_Fluc_ LS and Huh7_Rluc_ cells. (*Pf* BS sheet) Raw and transformed data for the dose-response effect of hit compounds on *P. falciparum* BS. (LS vs dual-stage sheet) List of compounds with LS-specific (IC_50_ *Pf* BS / IC_50_ *Pb* LS > 10) and dual-stage (IC_50_ *Pf* BS / IC_50_ *Pb* LS < 10) activity. Only compounds with a SI (CC_50_ Huh7 / IC_50_ *Pb* LS) > 10 were included.

